# An endolysosome membrane transformation process for engulfment of autophagosomes independently of ESCRT

**DOI:** 10.1101/2022.05.05.490766

**Authors:** Zhe Wu, Yufen Wang, Chuangpeng Wang, Fengping Liu, Yueguang Rong

**Affiliations:** School of Basic Medicine, Tongji Medical College, Huazhong University of Science and Technology, Wuhan, 430030, China; Cell Architecture Research Center, Huazhong University of Science and Technology, Wuhan, Hubei 430030, China

## Abstract

Endolysosome, usually regarded as the cellular recycling bin, digests materials from multiple origins. The materials from different origins are delivered to endolysosome through the vesicle fusion, kiss-and-run mechanisms, CMA or microautophagy. However, it remains unknown whether endolysosome can receive cargo via other ways. Here, we reported another endolysosomal process for receiving materials. In this, endolysosome receives materials by extending two arms which embraces autophagosomes and engulfs the autophagosomes ultimately, but not via the conventional fusion of autophagosomes with endolysosomes. We named this process as LEA (lysosome eats autophagosome) and the endolysosome arms as LF (endolysosome filopodia) provisionally. The endolysosomes with engulfed autophagosomes (LEA endolysosomes) get more autophagosomes via the fusion with other LEA endolysosomes. The engulfed autophagosomes are labelled by F-actin on their membranes and have ER protein Sec61β and peroxisome protein Pex14 inside their lumens, but mitochondria are excluded outside endolysosome. Our discovery of LEA and LF reveal an unidentified endolysosome transformation process which is responsible for receiving cargoes.

## Introduction

Endolysosome is usually regarded as a degradative organelle receiving lysosomal hydrolases from biosynthetic pathway turns over the materials from outside and inside of cells, such as endocytosis, phagocytosis, macroautophagy, microautophagy, CMA, MDV, mitochondria-lysosome fusion^1-3^. In endocytosis and phagocytosis, the engulfed materials from outside is delivered to endolysosome via early endosomes and late endosomes by vesicle fusion or kiss-and-run^1,2^. Intracellular material delivery to lysosomes is carried out via autophagosome-lysosome fusion^4,5^, MDV-lysosome fusio^6-8^ and mitochondria-lysosome fusion^9^. Lysosomes or vacuoles (the equivalent of lysosome in yeast) enclosed cytoplasmic cargoes, such as cytosolic proteins, lysosomal membrane proteins, ER, peroxisomes, nucleus and mitochondria via microautophagy dependently of the ESCRT complex^2,3^. However, whether endolysosomes receive cargoes via other modes remains unknown. Here, we found when endolysosome activity and mTOR activity are inhibited, the autophagosomes attach to endolysosomes and then endolysosomes transform to extend two arms out. The two arms embrace autophagosomes and ultimately engulfed autophagosomes possibly via a homotypic fusion process between the arms or between the arms and the endolysosome boundary membranes.

## Results

It is known that STX17, an autophagosomal SNARE for autophagosome-lysosome fusion, is only recruited to the outer membrane of sealed autophagosomes in wild type cells^10^. The GFP-tagged transmembrane domain of STX17 (GFP-STX17 TM) can also be recruited to the outer membrane of autophagosomes and translocates to autolysosome via autophagosome-lysosome fusion during autophagy^11^. But our previous report found that STX17 enters endolysosomes in a vesicle form upon EBSS and bafilomycin A1 (BFA) treatment^12^. In autophagosome formation deficient cells (FIP200 KO and ATG9A KO) the entrance of STX17 is blocked^12^, suggesting the STX17 signal inside endolysosome comes from autophagosomes. However, how these STX17 molecules enter lysosome remains unknown.

To further explore the mechanisms for the entrance of STX17, cells were treated with Torin2+ BFA, rapamycin+BFA, EBSS+chloroquine (CQ), Torin2+CQ, rapamycin+CQ, we found GFP-STX17 TM also enters lysosomal lumen upon these treatments **(Figure 1)**. These result suggest both endolysosome acidification and mTOR activity promotes the STX17 TM entrance into endolysosome.

**Figure 1.**
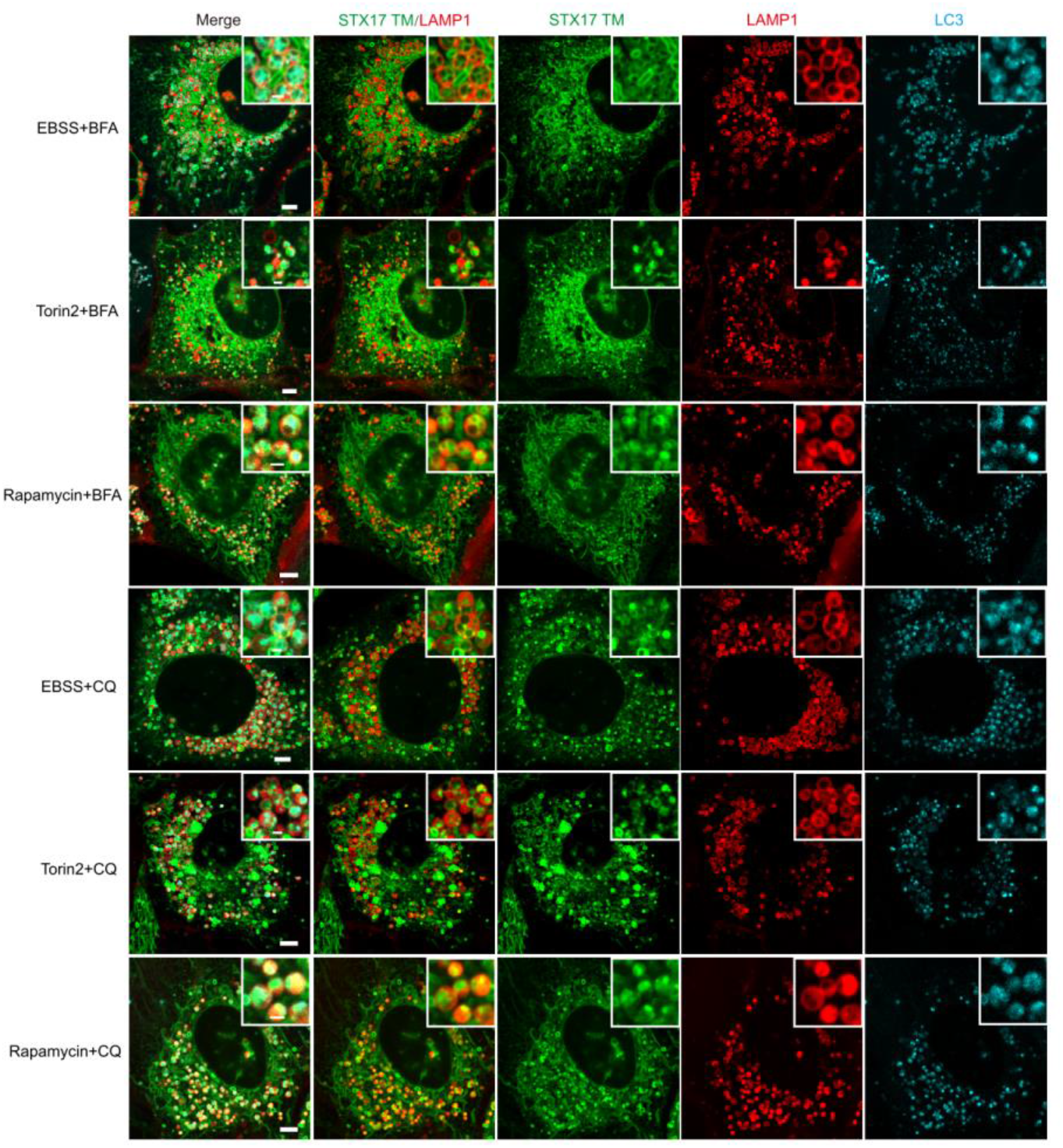
STX17 positive autophagosomes enter endolysosome upon acidification and mTOR inhibition. U20S cells stably expressing GFP-STX17 TM, LAMP1-mCherry and CFP-LC3 were treated with EBSS+BFA, Torin2+BFA, Rapamycin+BFA, EBSS+CQ, Torin2+CQ and Rapamycin+CQ for 5 hours, respectively and then cells were imaged by microscopy. Scale bar, 5μm. Inset scale bar, 1μm.

There are two possibility for the STX17 TM entrance into endolysosomes: 1. Some STX17 presents on inner membrane of autophagosomes. When autophagosomes fuse with endolysosomes, the STX17 positive autophagic bodies are released into the endolysosomal lumens. 2. Endolysosomes engulf autophagosomes by the other way.

We excluded the first possibility based on the following evidence: 1. It is well known that STX17 is only recruited to the outer membrane of autophagosomes in wild type cells^11^. Therefore, after autophagosome-lysosome fusion, the STX17 should present on the lysosomal membrane, but not in the lysosomal lumen. 2. Further, we observed the autophagosome-endolysosome fusion process under EBSS+BFA treatment by live cell imaging. The live cell images revealed that STX17 TM signal transferred from autophagosome membrane to endolysosome membrane via autophagosome-lysosome fusion, but not entered endolysosome lumen **(Figure 2A)**. 3. GFP tagged full length STX17 (GFP-STX17) blocks the autophagosome-lysosome fusion^13^, however, the GFP-STX17 still enters endolysosome lumen upon EBSS+BFA, Torin2+BFA, rapamycin+BFA, EBSS+CQ, Torin2+CQ, rapamycin+CQ **(Figure 3A)**. In addition, the STX17 entrance to endolysosome remains unaffected in STX17 and YKT6 double depletion cells **(Figure 3B)** in which autophagosome-lysosome fusion is completely blocked^14^. All these results suggest the STX17 entrance into endolysosomes independently of autophagosome-lysosome fusion.

**Figure 2.**
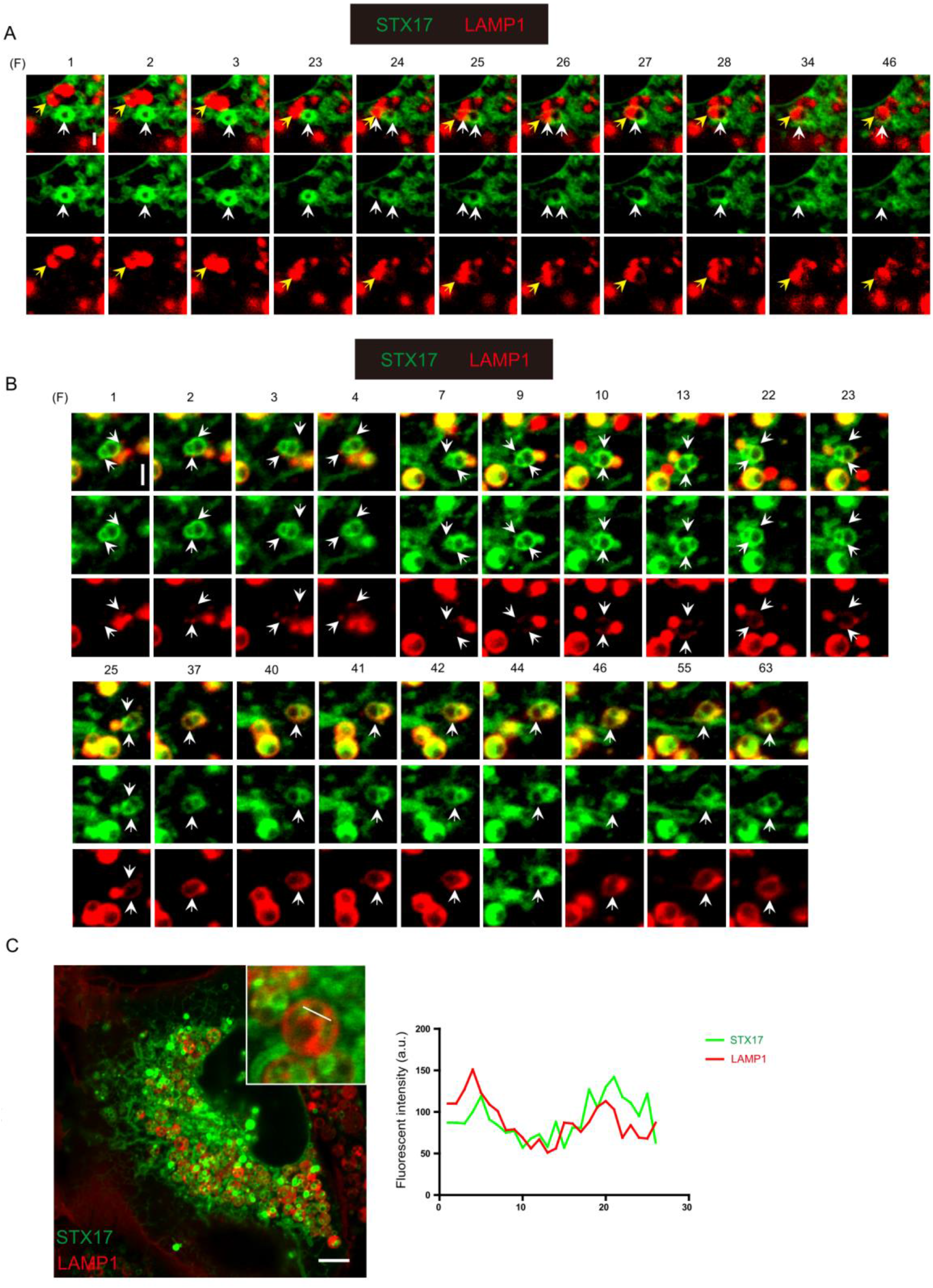
The STX17 positive autophagosomes fusion with lysosomes or engulfment by lysosomes upon acidification and mTOR inhibition. **A)** U20S cells stably expressing GFP-STX17 TM, LAMP1-mCherry were treated with EBSS+BFA for 2 hours and time-lapse images were taken by microscopy 18s/frame. The white and yellow arrows indicate the autophagosome and lysosome respectively during fusion. (F) represents frame. Scale bar, 1μm. **B)** U20S cells stably expressing GFP-STX17 TM, LAMP1-mCherry were treated with EBSS+BFA for 2 hours and time-lapse images were taken by microscopy 4.3s/frame. The white arrows indicate lysosomal extensions (LF). F represents frame. Scale bar, 1μm. **C)** U20S cells stably expressing GFP-STX17 TM, LAMP1-mCherry were treated with EBSS+BFA for 5 hours. Scale bar, 5μm. Right panel: fluorescent intensity of STX17 and LAMP1 on LEA autolysosomal membrane was analyzed.

**Figure 3.**
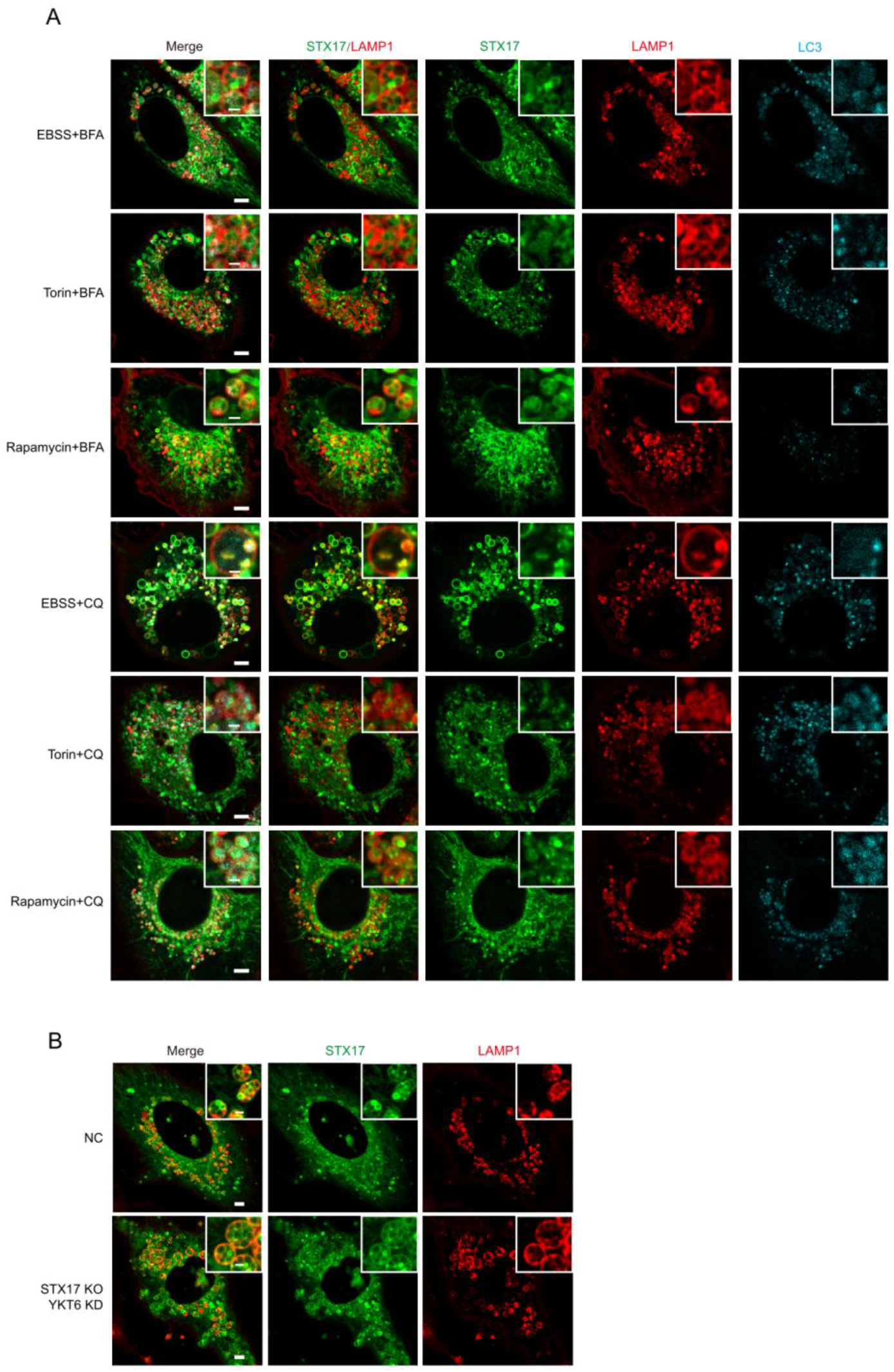
The engulfment of STX17 positive autophagosomes by lysosomes is independent of autophagosome-lysosome fusion. **A)** U20S cells stably expressing GFP-STX17, LAMP1-mCherry and CFP-LC3 were treated with EBSS+BFA, Torin2+BFA, Rapamycin+BFA, EBSS+CQ, Torin2+CQ and Rapamycin+CQ, respectively, for 5 hours and images were taken by microscopy. Scale bar, 5μm. Inset scale bar 1μm. **B)** STX17 KO U20S cells stably expressing GFP-STX17 TM, LAMP1-mCherry were transfected with non-targeting siRNA (NC) or siRNA against YKT6. Forty-eight hours after transfection, cells were starved with EBSS+BFA for 5 hours. Scale bar, 5μm. Inset scale bar 1μm.

Next, we tested the second possibility by live cell imaging to observe what happens to the endolysosome and the autophagosome upon EBSS+BFA treatment. From the live images we observed that an autophagosome attached to an endolysosome and then two arms with weaker LAMP1 signal extended from the endolysosome and gradually surrounded the autophagosome tightly; ultimately, these two arms encase the STX17 positive autophagosome into endolysosome lumen completely **(Figure 2B)**. This results suggest the STX17 positive autophagosome is engulfed into the endolysosome upon EBSS+BFA treatment. Consistently, the line profile analysis indicated that the LAMP1 signal is aligned with STX17 signal on the autophagosome membrane in endolysosomal lumen **(Figure 2C)**. Since this process has not been discovered in mammalian cells, therefore, we named this process as LEA (endolysosome eats autophagosome) and the transformation of the endolysosome as FL (endolysosome filopodia) for abbreviation provisionally.

We often observed several STX17 positive vesicles inside one endolysosome upon EBSS+BFA treatment, therefore time laps images were taken to observe how this happens. Interestingly, we found two endolysosomes with a STX17 vesicle inside respectively, attached together and fused to form a bigger endolysosome with two STX17 vesicles. After this fusion, the resultant endolysosome fused again with the other endolysosome nearby with several STX17 vesicles to become a bigger endolysosome with more STX17 vesicles **(Figure 4)**. This live cell images suggest the endolysosomes after LEA can gain more STX17 vesicles via endolysosome-endolysosome fusion.

**Figure 4.**
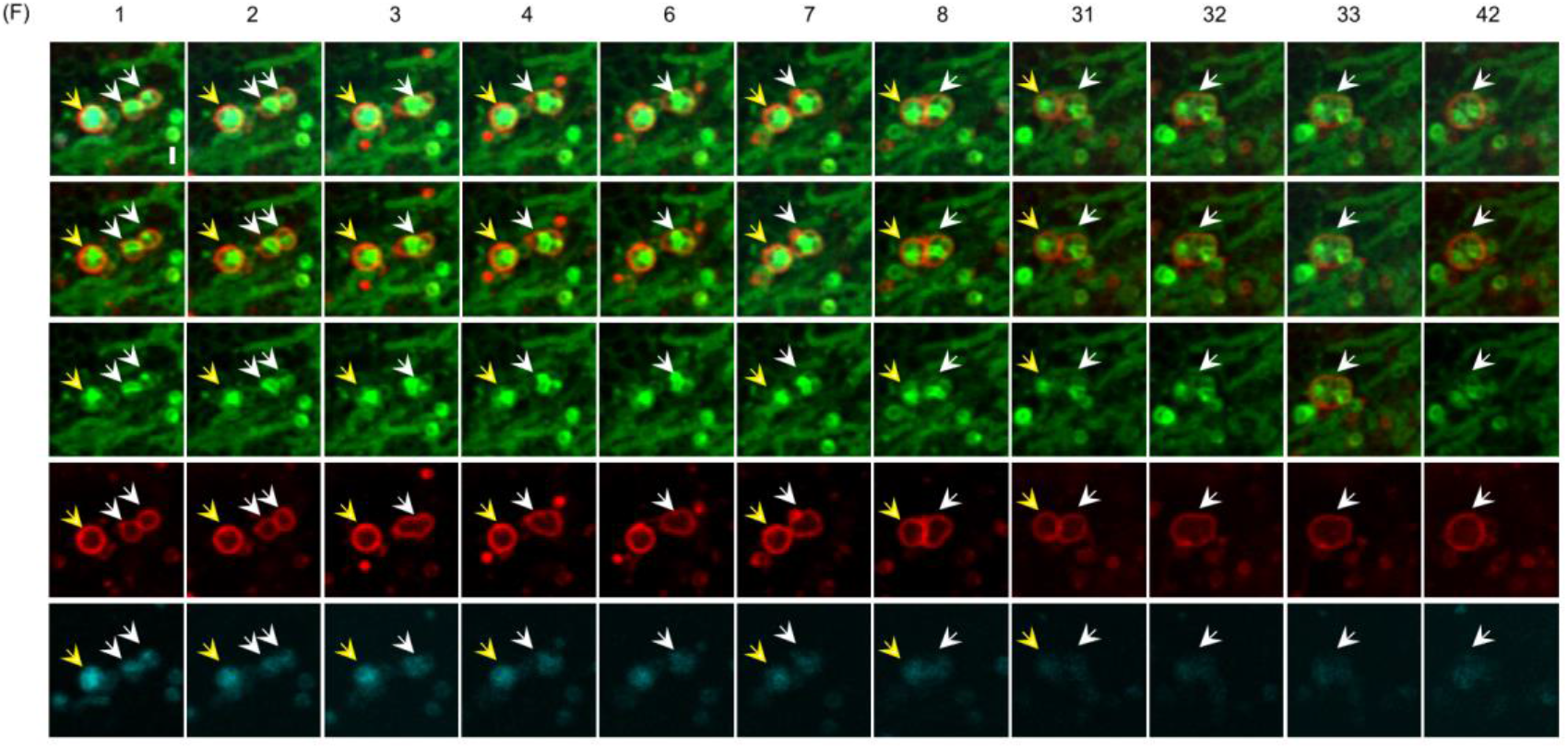
The LEA autolysosomes fuse each other. U20S cells stably expressing GFP-STX17 TM, LAMP1-mCherry and CFP-LC3 were treated with EBSS+BFA for 2 hours and time-lapse images were taken by microscopy 20s/frame. The white and yellow arrows indicate the different LEA autolysosomes. (F) represents frame. Scale bar, 1μm.

To further characterize this process, we expressed lifeact (which labels the F-actin), Sec61β(the ER marker), Pex16 (peroxisome marker), TOM7 (mitochondria marker) in LEA occurring cells. We found the membrane of STX17 positive autophagosomes inside endolysosomes the lifeact positive **(Figure 5A)**, which may derive from the endolysosome membrane attached actin^15,16^. In line with the previous report that actin assembles inside autophagosome precursors^17^, we also observed some dim lifeact signals inside these autophagosomes. The lumen of these STX17 positive autophagosomes have Sec61β puncta inside **(Figure 5B)**, consistent with the autophagosome cradle model^18^. We also found the Pex16 positive puncta enclosed inside the LEA autophagosomes **(Figure 5C)**. Interestingly, we rarely observed mitochondria were enclosed into the STX17 autophagosomes inside endolysosomes or into endolysosomes directly during LEA. Even when the cells were treated with oligomycin and antimycin A (OA) which leads to mitochondria fragmentation **(Figure 6)**. This suggests the treatment here triggers LEA cannot induce the engulfment of mitochondria by endolysosomes.

**Figure 5.**
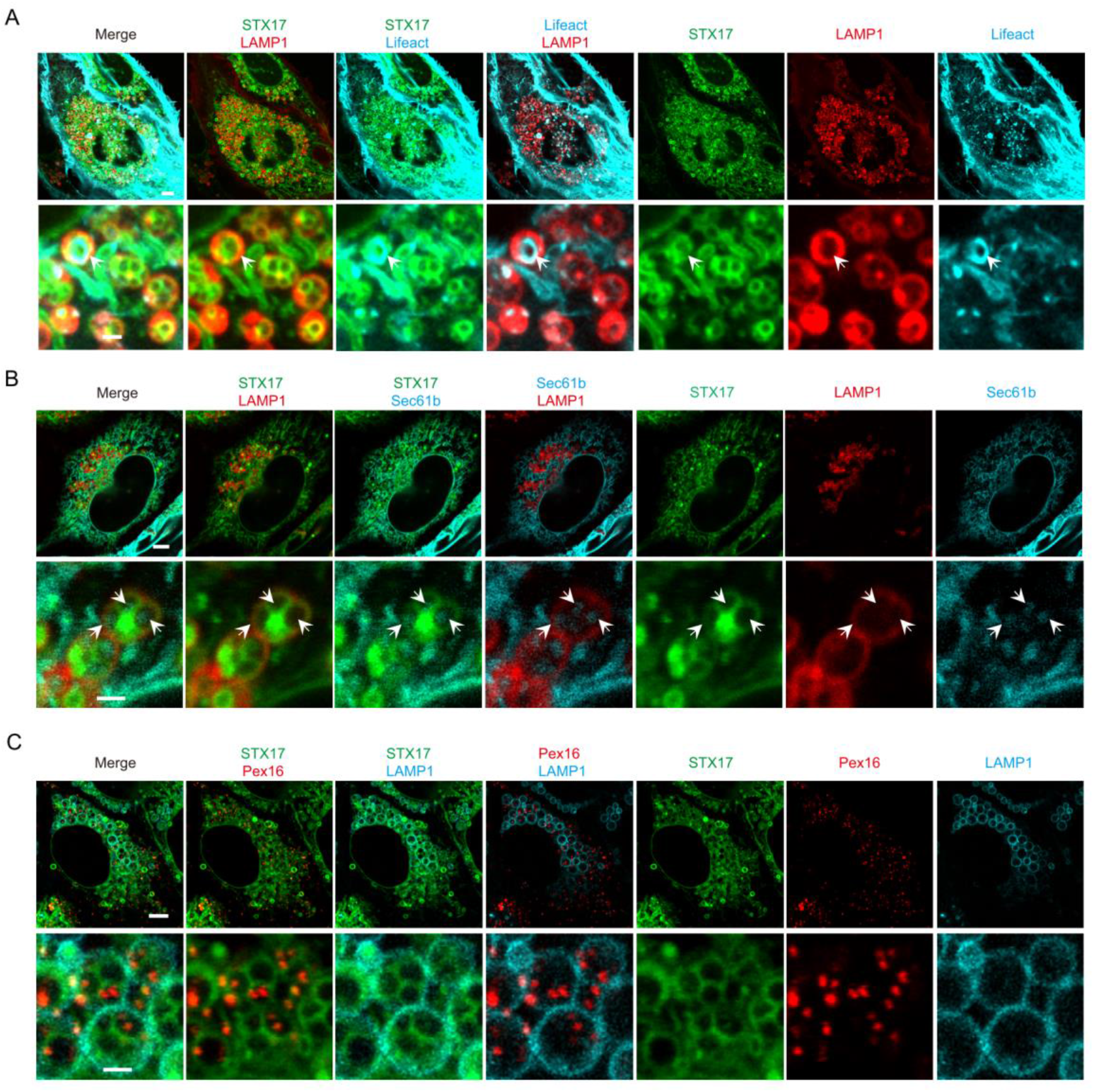
The localization of actin, Sec61βand Pex16 in LEA autolysosomes. **A)** U20S cells stably expressing GFP-STX17 TM, LAMP1-mCherry and lifeact-CFP were treated with EBSS+BFA for 5 hours and images were taken by microscopy. Scale bar 5μm. Inset scale bar 1μm. **B)** U20S cells stably expressing GFP-STX17 TM, LAMP1-mCherry and CFP-Sec61βwere treated with EBSS+BFA for 5 hours. Scale bar 5μm. Inset scale bar 1μm. **C)** U20S cells stably expressing GFP-STX17 TM, Pex16-mKATE2 and LAMP1-CFP were treated with EBSS+BFA for 5 hours. Scale bar 5μm. Inset scale bar 1μm.

**Figure 6.**
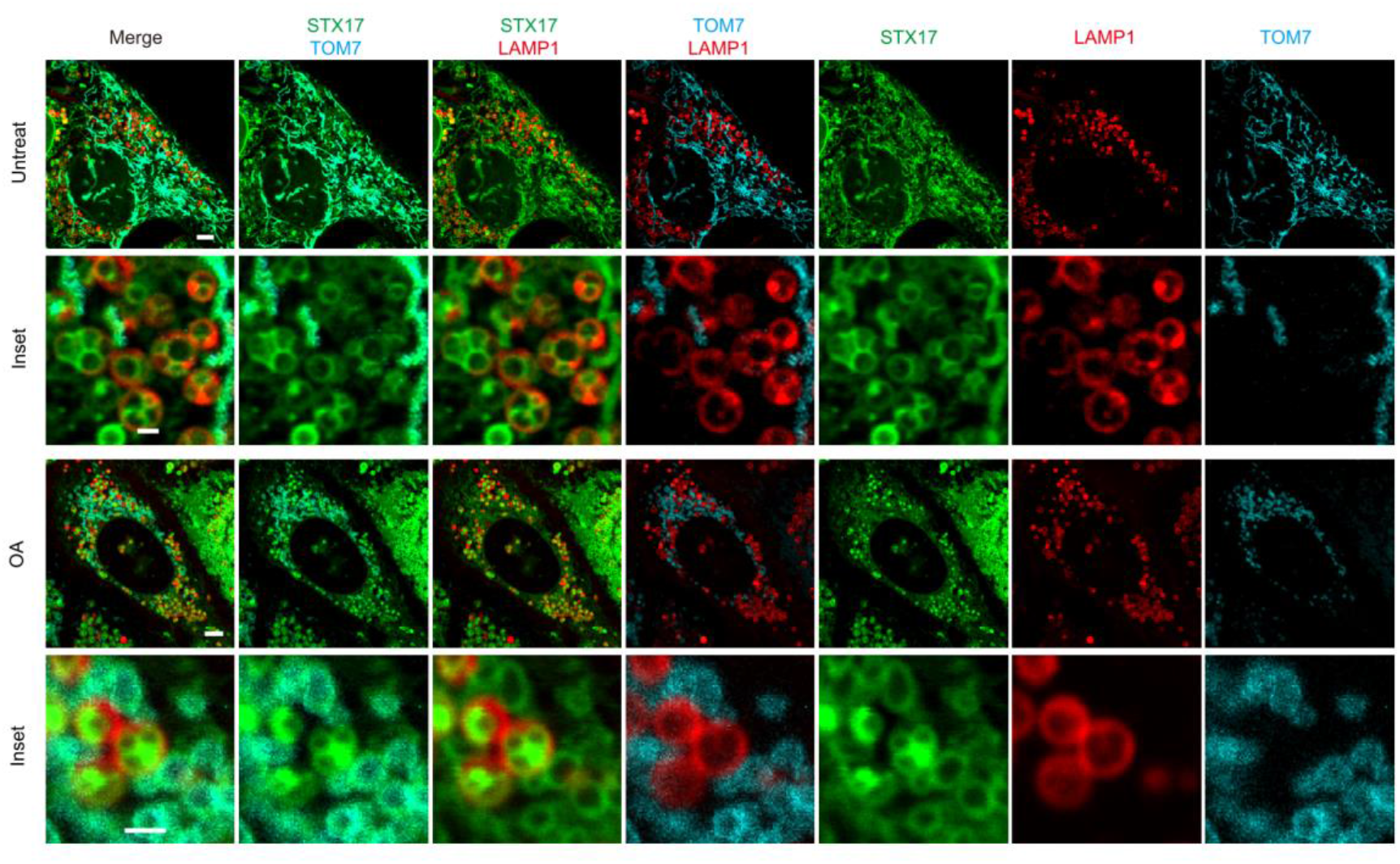
LEA autolysosomes exclude mitochondria. U20S cells stably expressing GFP-STX17 TM, LAMP1-mCherry and CFP-Tom7 were treated with OA (oligomycin and antimycin A) and EBSS+BFA for 5 hours and images were taken by microscopy. Scale bar 5μm. Inset scale bar 1μm.

Since the ESCRT complex has been involved in many invagination processes on endosomes and lysosomes^3^, we examined whether ESCRT is involved in LEA. Unexpectedly, we found LEA is not affected in cells knockdown of the subunits of ESCRT-0, ESCRT-I, ESCRT-II, ESCRT-III, VPS4A/B and ESCRT related genes ALIX and UBPY **(Figure 7)**. These results suggest the ESCRT complex is dispensable for LEA.

**Figure 7.**
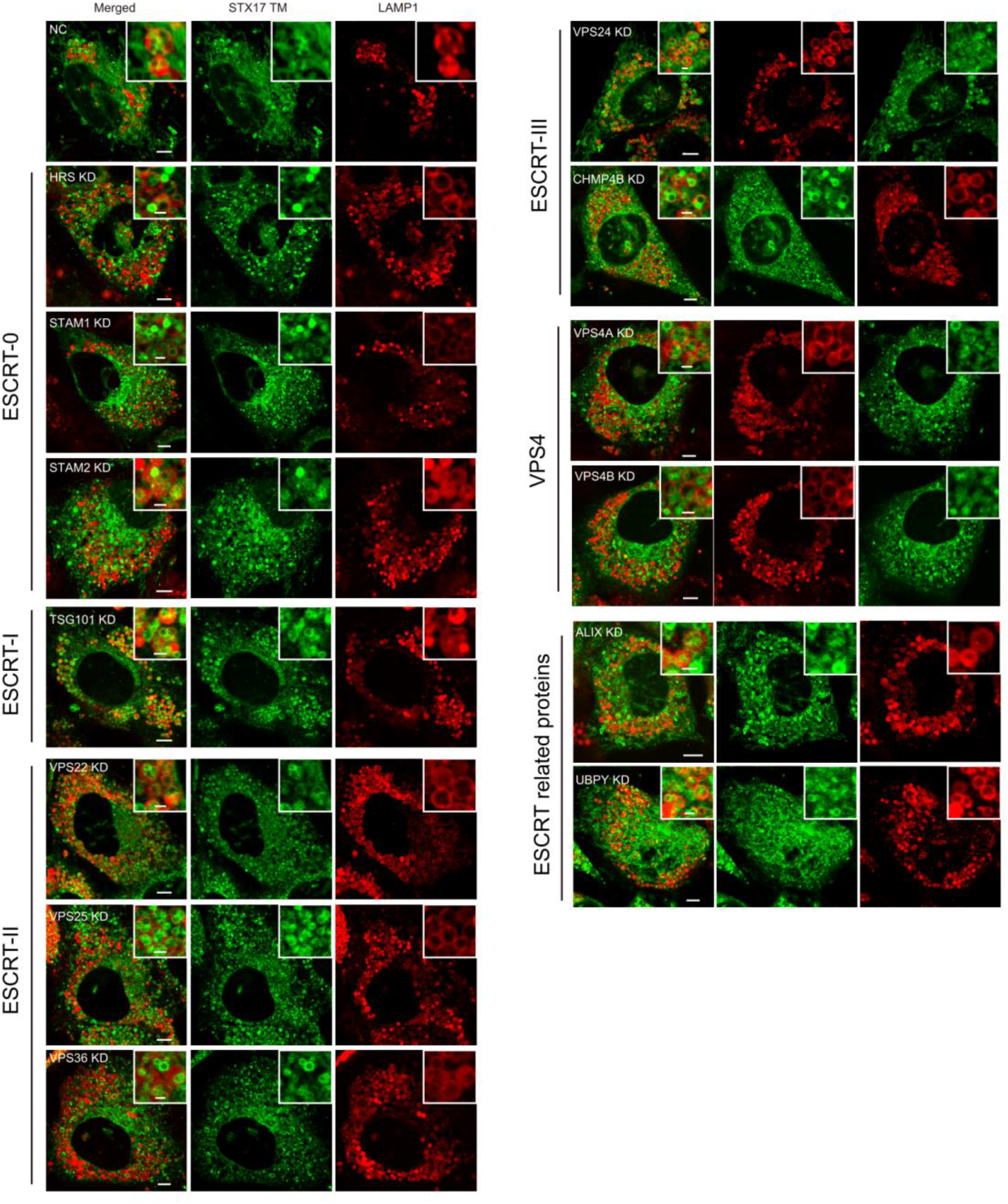
ESCRT is dispensable for LEA. U20S cells stably expressing GFP-STX17 TM, LAMP1-mCherry were transfected with non-targeting siRNA (NC) or siRNA against the individual ESCRT genes or ESCRT related genes. Forty-eight hours after transfection, cells were starved with EBSS+BFA for 5 hours. Scale bar, 5μm. Inset scale bar 1μm.

## Discussion

Here, we discovered an unidentified endolysosome transformation process LEA (endolysosomes eat autophagosomes), in which LF (endolysosome filopodia) from endolysosome engulfs autophagosomes, when endolysosome acidification and mTOR activity is inhibited. Both of autophagosome-lysosome fusion and ESCRT are dispensable for this process.

Many membrane invagination processes on endosomes and lysosomes/vacuoles, such as microautophagy on endosomes and lysosomes, are mediated by ESCRT^3,19^. But we found ESCRT is dispensable for LEA, even in some ESCRT deficient cells LEA efficiency is increased. Further, the diameter of intraluminal vesicles generated by ESCRT in MVB is around 25 nm (50 nm in human cells) and 200nm-500nm in lysosomes/vacuoles^3,19^. Here, the vesicles in LEA is usually around 1μm. Given the LEA is executed in an ESCRT independent manner and the bigger size of its intraluminal vesicles, we think LEA is a different process from microautophagy on endosomes and lysosomes and therefore the detailed mechanism also remains to be investigated.

We found the endolysosome membrane extrusion in LEA, we named it as endolysosome filopodia (LF), for enclosing cargos, which is a dramatic difference from the known endolysosome processes. For microautophagy either in yeast or mammalian cells, the lysosomes/vacuoles take up cargoes by invagination. This is possibly because vacuoles are relative bigger than endolysosomes, and therefore vacuoles invaginate cargos inside, while in LEA the mammalian endolysosomes are very small, it is impossible for endolysosomes to invaginate the large cargos. Although mammalian endolysosomes also engulf smaller cargos, such as ER, dependently of LC3 interaction with Sec62 and ESCRT^3,20^. Probably, mammalian endolysosomes evolve to from membrane extrusions for engulfment of bigger cargos. The mechanisms for membrane extrusion should be different from the ESCRT dependent invagination.

Further, LF displays very weak LAMP1 signal intensity which is not easy to be observed, suggesting this portion of membrane is different from endolysosome boundary membrane in properties, which enriches the lysosomal abundant membrane proteins such as LAMP1. It suggests endolysosomes segregate a specific portion of membrane for LEA. In micro-lipophagy, lipid droplets are invaginated in L0 domains (liquid-ordered domains)^21-23^. Nuclear pore complexes (NPCs) were excluded from NV junctions^24^. In micro-pexophagy, the MIPA as an independent and separate membrane covers the invaginated peroxisomes on the opposite side of vacuoles^25^. Therefore, it seems a common phenomenon for endolysosomes/vacuoles to separate a membrane portion for special functions, as it occurs in microautophagy or LEA. According to our results, the MIPA-like structures for micro-pexophagy sealing is not observed in LEA. Likely the last step for LEA is executed by the homotypic fusion of LF fusion with endolysosome boundary membrane, thus fusion machineries may be involved.

LEA only engulfs the STX17 positive autophagosomes and autophagosome enclosed substrates, but exclude mitochondria, suggesting LEA recognize cargo with specificity, rather than randomly. But how lysosomes achieve this specificity remains unknown and deserved to be investigated in future. Although here endolysosomes engulfs autophagosomes only, the endolysosome transformation process possibly is also used for receiving other cargoes under other conditions. Therefore, the cargoes engulfed by endolysosomes via LF remains to be identified under different conditions.

In FIP200 and ATG9A KO cells, the LEA is completely diminished. Given the lack of autophagosomes in autophagosome formation deficient cells, likely the block of LEA is an indirect effect from the inhibition of autophagosome formation. However, we cannot exclude the possibility that these genes also have a direct role in LEA.

Upon lysosome acidification and mTOR inhibition, the time laps images revealed that the some STX17 positive autophagosomes also fuse with lysosomes and displaying STX17 signaling on autolysosomal membranes. It suggests macroautophagy occurs concurrently with LEA, though with a low activity compared with LEA.

## Method

### Live cell imaging

Cells were placed on a four-chambered cover glass (In vitro scientific, syD35-20-1-N) 1 day before observation. During live imaging, the culture dish was mounted on an inverted microscope (Olympus, FV3000) to maintain incubation conditions at 37? and 5% CO2 using a Plan Apochromat N 100×/1.70 oil. Images or videos were recorded using confocal laser microscope system, then further processed and analysed using ImageJ.

### Contributions

The author ordering of Zhe Wu, Yufen Wang, Chuangpeng Wang, and Fengping Liu is unrelated to their contributions.

